# DeepInsight-FS: Selecting features for non-image data using convolutional neural network

**DOI:** 10.1101/2020.09.17.301515

**Authors:** Alok Sharma, Artem Lysenko, Keith A Boroevich, Edwin Vans, Tatsuhiko Tsunoda

## Abstract

Identifying smaller element or gene subsets from biological or other data types is an essential step in discovering underlying mechanisms. Statistical machine learning methods have played a key role in revealing gene subsets. However, growing data complexity is pushing the limits of these techniques. A review of the recent literature shows that arranging elements by similarity in image-form for a convolutional neural network (CNN) improves classification performance over treating them individually. Expanding on this, here we show a pipeline, DeepInsight-FS, to uncover gene subsets of clinical relevance. DeepInsight-FS converts non-image samples into image-form and performs element selection via CNN. To our knowledge, this is the first approach to employ CNN for element or gene selection on non-image data. A real world application of DeepInsight-FS to publicly available cancer data identified gene sets with significant overlap to several cancer-associated pathways suggesting the potential of this method to discover biomedically meaningful connections.

## Introduction

Traditional machine learning (ML) algorithms (such as support vector machines ^1^, random forest ^2^, and logistic regression ^3^) are dominantly applied in classification and feature selection of non-image data. Primarily, a *d* × 1 column vector is supplied to an ML algorithm to find a smaller set of features and/or to identify it into one of the defined categories. In the medical field, ever-increasing data complexity is pushing the limits of ML algorithms to extract relevant information for phenotype identification related to disease diagnosis and analysis. In this respect, the selection of a small subset of critical elements or genes from a larger set of elements has become a critical step. The element or gene selection (also known as feature selection) problem is not limited to genomic data analysis but is an important process in many areas of research. The reliability of ML algorithms to find a subset of genes is mostly determined by the feature selection, feature extraction and classification steps.

On the other hand, convolutional neural networks (CNNs) are a class of deep learning architectures that have shown promising results and gained widespread attention concerning image data ^4-14^. CNN takes an input image (a *p* × *q* feature matrix) and through its hidden layers conducts feature extraction and classification. One of the key advantages of CNN is their high efficiency, i.e. fewer samples and less training time are needed to achieve good levels of performance. This led to their high popularity in a myriad of cutting-edge commercial applications (e.g. driverless cars). CNN has several advantages: it derives features from raw elements and does not require any additional feature extraction technique, it finds higher-order image statistics and nonlinear correlations, it requires less convolutional neurons as it processes data for its receptive fields (or small sub-area), allowing a deeper network with fewer parameters, and its receptive fields share the weights and biases, reducing the memory footprint ^15^. As input, CNN takes an image. In a local region, an image is comprised of spatially-coherent pixels; i.e., similar information is shared by the pixels near each other. If pixels are arbitrarily arranged, then their placement can negatively impact the performance of the feature extraction and classification process. Consequently, the order of the adjacent pixels of an image in CNN is not independent. This is different from ML techniques as the latter are not impacted by a change in the order of the elements. Luckily, for CNNs, the images utilized are usually a representation of physical entities and, therefore, do not require pixel rearrangement, as the lenses of camera correctly pass the appropriate light shades of animate or inanimate objects to the pixels. Nonetheless, so far the applications of CNNs have been limited for non-image data (e.g. transcriptomic, genomic, single-cell RNA-seq, banking, finance, etc.).

One possible way CNN can process a non-image sample is by first converting it to an image sample considering spatially coherent pixels in its local regions. In this respect, the DeepInsight approach ^16^, pioneered by incorporating element arrangement through t-SNE mappings, feature extraction, and classification steps, was proposed. The element arrangement is carried out by positioning elements or genes on a 2D pixel frame based on their relative similarities, followed by the mapping of element values on these locations. This approach ubiquitously transforms non-image samples to images for CNNs. To our knowledge, it was the first approach to convert various kinds of non-image data to image forms for the application of CNN architecture. Lyu and Haque ^17^ previously applied CNN to RNA-seq data. They first carried out gene selection based on variance across all samples and then construed the remaining genes as images depending on chromosome location. Since this approach involves information about the location of chromosomes, it cannot be applied to other types of datasets. However, it was an essential contribution in this direction. Recently, Buturivić and Miljković ^18^ introduced an approach where tabular data for CNN (with ResNet architecture) is used by transforming rows of tabular data as an image filter, and then by applying it to a fixed-base image. They applied their approach to gene expression data derived from blood samples of patients with bacterial or viral infections and showed promising results over many ML algorithms. Kanber ^19^ applied the DeepInsight approach to the sparse data of the MINST database of 70k samples and showed it had superior performance than a state-of-the-art machine learning (random forest) method.

To date there are very few studies about how to perform feature selection by CNN for non-image samples, such as finding a subset of genes. In this work, we focus on developing a methodology to show that gene selection can be done using CNNs. The hidden layers of CNNs can reveal complex mechanisms (such as pathways) for non-image samples. The same methodology can be extended to other kinds of non-image cases and is not restricted to genomic or transcriptomic data. To this end, the proposed DeepInsight-Feature Selection algorithm, abbreviated as DeepInsight-FS, was developed. The DeepInsight-FS approach encompasses four main steps: element arrangement, feature selection, feature extraction and classification (please see Supplement File 1 for the definition about these terms). Its components are briefly summarized here. The pipeline of DeepInsight-FS is shown in Figure 1. The steps of DeepInsight-FS are discussed in the following section.

**Figure 1.**
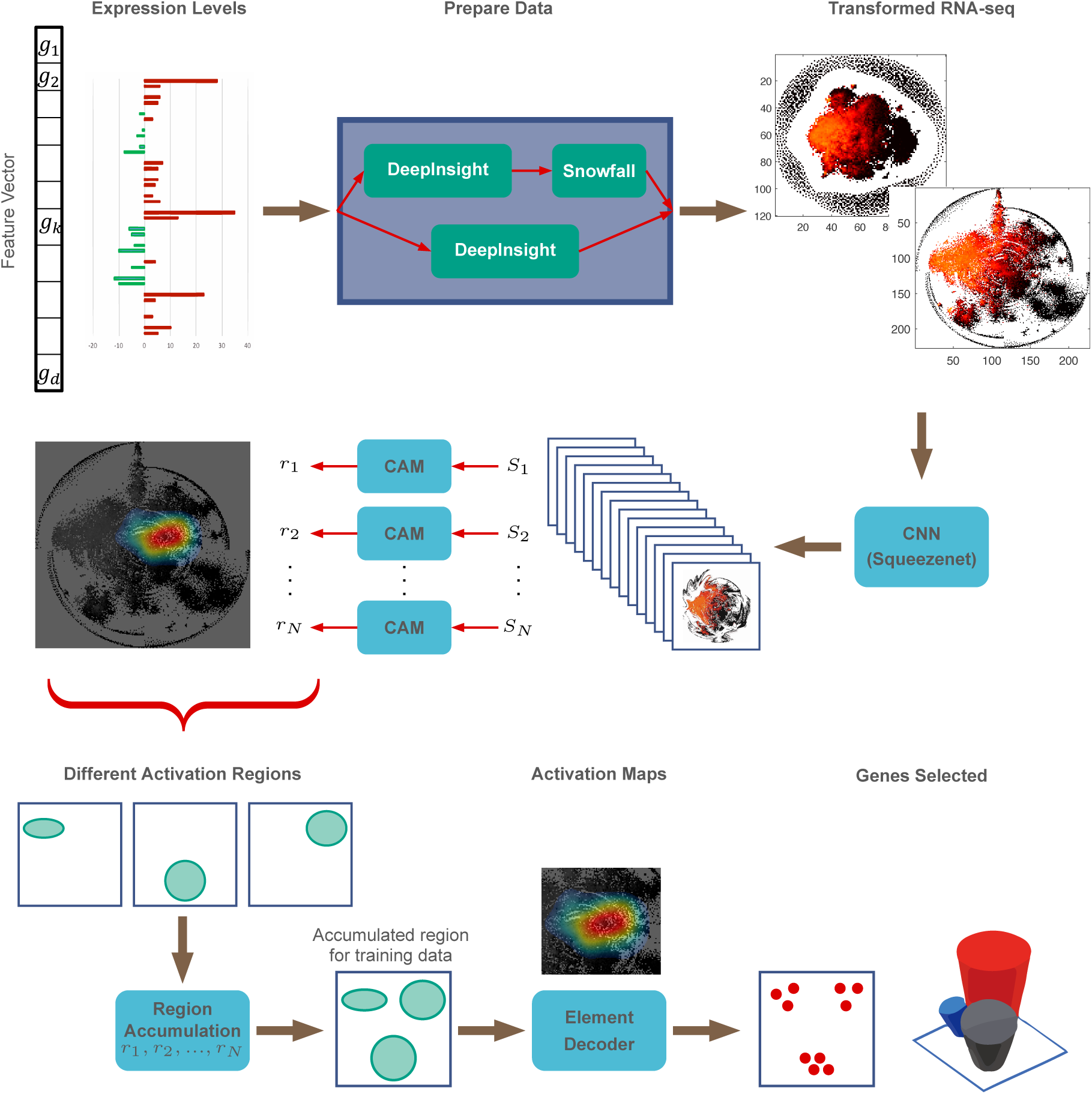
An overall procedure of selecting features using convolutional neural network. The model takes non-image data, e.g. transcriptomic or RNA-seq data, and finds a subset of genes or elements via CNN. The transformation of non-image samples to image samples is done following the element arrangement step of the DeepInsight model ^16^. Since the number of elements or dimensionality of data can be high, a snow-fall compression algorithm (details are discussed later in the manuscript) was developed to fit more genes or elements in a given pixel-frame to enable every possible gene to be part of the selection. The image samples obtained from the DeepInsight and snow-fall algorithms are submitted to the CNN model (using SqueezeNet architecture ^20^). The hyperparameters of CNN are tuned with the training set and the model’s fitness is evaluated using the validation set. An independent test set was kept aside during the training phase of the model. The feature extraction and classification are performed by CNN. The feature selection is performed collectively by the class activation maps (CAM) ^21^, region accumulation, and element decoder. CAM are used to find activations of each sample. The activations for individual samples are integrated to find active regions for a class or category at the region accumulation step. The accumulated regions (for one or all classes) define pixel locations of interest for categorization of samples. These selected pixels are decoded to provide a subset of elements at the element decoder step. If the number of selected genes is higher than the desired number of genes, then the whole procedure can be executed again with the selected genes as input to find further subsets of genes. Repeating these steps will reduce the number of selected genes.

## Results

The results are produced by following an overall procedure of DeepInsight-FS (Figure1). The model takes non-image data, e.g. transcriptomic or RNA-seq data, and finds a subset of genes or elements via CNN. The model carries the following steps, image transformation by DeepInsight, snow-fall compression to enable more elements in a pixel-frame, classification via SqueezeNet model of CNN architecture, finding activation maps by CAM model, discover the overall activated regions for a category or cancer type via the region accumulation step, and decoding gene subsets by the element decoder procedure.

First, we compared the number of selected genes and classification performance for cancer type identification of different machine learning algorithms (Table 1). Lasso gave 1018 non-zero coefficients and as a result it discarded all other elements or genes. For ANOVA and variable genes method, 1000 genes are extracted.

**Table 1:** Classification accuracy and gene selection using machine learning and DeepInsight-FS algorithms.

| Machine learning methods | #genes selected | Classification accuracy |
| --- | --- | --- |
| ANOVA+RF | 1000 | 94% |
| Lasso+RF | 1018 | 96% |
| Variable genes+RF | 1000 | 92% |
| Logistic regression+RF | 687 | 97% |
| DeepInsight-FS | 1806 | 98% |

These algorithms, with the exception of logistic regression, provide one subset of genes for all the categories. However, in some cases pairwise analysis is possible (e.g. by using post-hoc Tuckey’s test). In such case, a total of 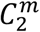 subsets are to be generated, where *C* is the combination function and *m* is the number of categories. On the other hand, logistic regression provides a separate model for each cancer type. This way, it is possible to find a subset of genes corresponding to a particular cancer type. We obtained 180 genes per cancer type, which gives 180 × 10 = 1800 genes for all the 10 cancer types out of which 687 genes are unique. Since the test classification accuracy for logistic regression is the highest among the existing methods studied in this paper, we consider it as the benchmark method for enrichment analysis. Moreover, other than DeepInsight-FS and logistic regression methods, no other methods gave gene sets respective to each of cancer types.

DeepInsight-FS produced 98% classification accuracy on the independent test set. The four distances Chebychev, correlation, cosine and hamming are used for DeepInsight-FS to obtain a subset of 5228 genes (see Tables S1.1 - S1.3 for hyperparameter details, Figure S1.2 for an illustration of corresponding activations and Figure S1.3 for gene subsets per cancer type). This subset of genes was further processed with DeepInsight-FS using hamming distance to achieve 1806 genes. The details about DeepInsight-FS execution and results of various stages involved to achieve 1806 genes can be found in Supplement File 1.

DeepInsight-FS is also capable of finding different gene subsets belonging to different cancer types. We applied enrichment analysis on 1806 genes obtained belonging to 10 cancer types. The number of genes for in each cancer type was in the range between 930 and 1450 (see Figure S2.1 in Supplement File 2 for genes arrangement in cancer types). DeepInsight-FS has more consistently identified a common set of genes which were selected for all cancer types (Figure 2a and Figure S2.1; Figure S2.2 for performance evaluation of all the algorithms; and, Figure S2.3 for overlap of gene subsets among all the methods including gene annotation). Interestingly, in addition to this overall trend, logistic regression and DeepInsight-FS appear to preferentially select the opposite sets of genes relative to each other. However, DeepInsight-FS was far better in recovering significantly enriched cancer-relevant pathways (Figure 2b). This suggests both that the proposed algorithm is better at discovering biologically coherent groupings of genes and that most of these grouping also appear to be highly relevant to cancer-associated processes. Notably, sets of genes from DeepInsight-FS are also depleted for housekeeping genes (Figure 2a). As housekeeping genes usually tend have stable expression across all cell types and tissues, this result likely an indication that fewer false positive genes, which are unlikely to be of interest, were chosen.

**Figure 2:**
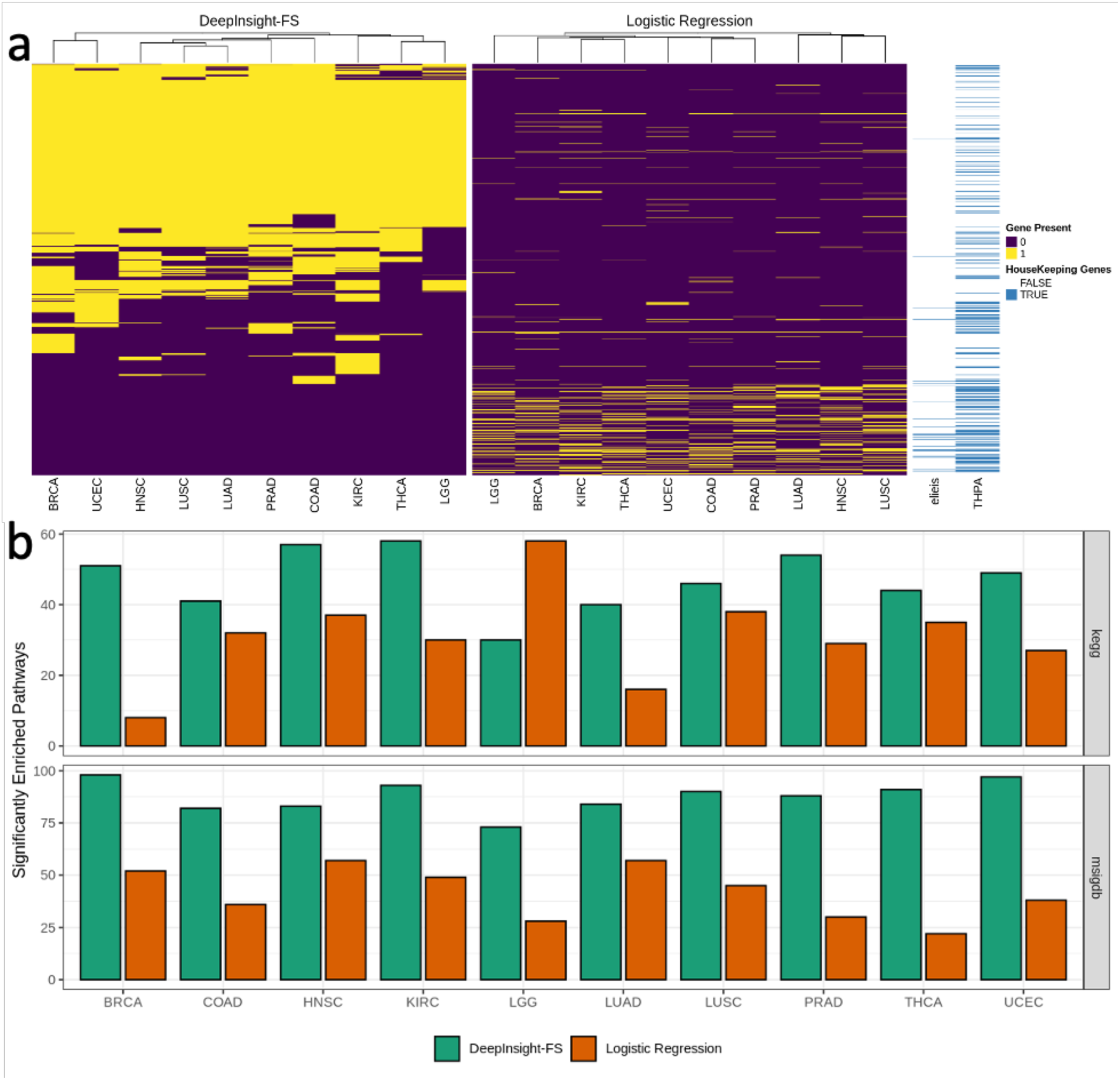
a) Commonality of gene selected by DeepInsight-FS (DI-FS) and logistic regression (LR) techniques. Each row represents a gene and yellow denotes that gene was selected for that cancer type (column). Housekeeping genes, as identified by two annotation sources, are shown in blue. b) Enrichment analysis in KEGG and MSigDB gene sets. The number of significantly enriched cancer sets for genes selected by DI-FS (green) and LR (orange) for each of the ten cancer types. DI-FS produced higher counts than LR for MSigDB for all the cancer types, and all but one (LGG) for KEGG.

## Discussion

The development of DeepInsight-FS opens the possibility of applying CNNs for feature selection problems, augmenting the power of CNNs for many real-life applications. The DeepInsight-FS method conducts element arrangement, feature selection, feature extraction and classification operations. The element arrangement step places similar elements together on a pixel-frame allowing meaningful image conversion of non-image samples. The selection process through activation points provides a subset of useful elements or genes. Layers of CNN perform the feature extraction and classification tasks.

The performance evaluation of DeepInsight-FS is gauged against finding a subset of genes through enriched pathways and classification accuracy on the independent test set. As expected, promising results were obtained when compared with the state-of-the-art machine learning techniques. The enrichment counts for DeepInsight-FS were reasonably sound compared to the benchmarked machine learning technique. The classification accuracy on the independent test set was 98%, which is also close to the machine learning technique. This shows that deep learning architectures have a possibility to provide solutions for biomarker discovery, genomic analysis for a variety of input samples ranging from RNA-seq to various omics data. In general, this method is suitable for applications where given data is in non-image form.

Feature selection results indicated that DeepInsight-FS was much better than alternative algorithms at recovering biologically-meaningful groups of genes that were relevant to classes (phenotypes) of interest. Therefore, the algorithm is likely to have great utility for tasks like prioritization of diagnostical signatures and interpretation of complex multi-omics data.

It is now generally accepted that in order to fully understand the dynamic quintessential complexity of cancer, it is essential to profile and collect data using a wide array of possible methods. As a result, a typical study commonly needs to consider wide array of possible data types, possibly including images, clinical records, as well as protein-coding and non-coding RNA expression profiles and full-genome sequencing. Understanding mechanisms and discovery of clinically-relevant subtypes frequently requires appropriate aligning of these different types of data and, most often, subsequent use of machine learning algorithms to identify meaningful patterns. Among current generation of approaches, deep learning-based methods so far offer the greatest flexibility and can potentially allow fuller automation and more comprehensive integration of these different types. In principle, by combining different specialized encoder layers any types of data can be converted into a set of inter-compatible embeddings and used within the same machine learning system. This innate flexibility is difficult to match using other methods and is particularly valuable for mining complex multi-modal datasets.

In terms of specific pathways, DeepInsightFS has consistently identified genes belonging to “Extracelllular matrix organisation” pathway (Reactome: R-HSA-1474244), which was top group for all cancer types – both by number of genes and enrichment significance (see Supplement File 3 for top pathways). The importance of this pathway is in line with current understanding with its wider biological role, where it is known to be both highly diverse in its expression across different tissues and subject to dysregulation in cancer ^36^. Another two pathways that were most-enriched across all cancer types were “Signaling by Receptor Tyrosine Kinases” (R-HSA-9006934) and “GPCR ligand binding” (R-HSA-500792). Tyrosine kinases mediate both cell proliferation and apoptosis process and their importance for multiple types of cancers is relatively well-studied ^37^. Interestingly, although it was noted that different GPCRs are expressed in different cancer types and some of them may be suitable for use as biomarkers ^38^, overall roles of specific genes in cancer is not very well understood and ligand of many GPCRs still have not been identified ^39^. Further analysis of these DeepInsight-FS results, like holistic analysis of which parts of specific pathways were found to be important for different types, may help to explain key cancer type-specific differences, and allow to connect the GPCR signalling to other more well-understood mechanisms of oncogenesis.

Good correspondence of identified gene set with known gold-standard cancer associated pathways is particularly promising as a means of interpreting the deep learning results from a biological perspective. Despite many recent advances, deep learning still commonly has a reputation for generating high-quality but ultimately “black box” models, where discovering how an algorithm arrived at a particular conclusion is very challenging. However, given that structures of key cancer pathways are very well understood, their high degree of overlap with DeepInsight-FS results could be crucial for correctly contextualizing the importance of individual genes and offer possible explanations for why they were selected by the algorithm.

## Materials and Methods

### DeepInsight-FS: Feature selection for non-image data using CNN

This section defines the proposed DeepInsight-FS methodology. The constituents of the model are 1) image transformation by DeepInsight, 2) snow-fall compression to enable more elements in a pixel-frame, 3) SqueezeNet model of CNN architecture, 4) CAM model to find activation maps, 5) region accumulation to obtain overall activated regions for a category or dataset, and 6) element decoder to decode genes from active regions. These steps are discussed as follows.

### DeepInsight: non-image to image conversion for CNN

DeepInsight transforms a non-image sample to a well-organized image form by effectively arranging elements while considering neighborhood information. The feature extraction and classification tasks are done by CNN. DeepInsight integrates three steps: 1) element arrangement, 2) feature extraction and 3) classification. This cumulative approach of element arrangement can be useful in uncovering hidden mechanisms (e.g. pathways). This way the relative importance of features towards a target can be understood better. An input feature vector is transformed to a feature matrix using t-SNE ^22^, kernel PCA ^23^ or UMAP^24^, then the smallest rectangle containing all the elements is found using the convex hull algorithm. A necessary rotation is performed to align the image, and then Cartesian coordinates are converted to pixel coordinates. After that, mapping of element values on pixel locations are performed to construct an image of a feature vector. The details of image transformation have been previously described ^16^.

### Snow-fall Compression Algorithm

If the dimensionality of a sample with *d* elements, *x ∈* ℝ^*d*^, is very large, then it becomes very difficult to place all the elements in a given pixel frame of size *m* × *n*. Therefore, the question is, how to compress, such that all the elements can be arranged in the same pixel size, while maintaining the data topology. There are two ways of performing compression, lossy compression and lossless compression. In lossy compression, two or more elements can overlap, i.e., these elements will have an identical pixel location. In this case, the values of elements are averaged at that particular pixel location. On the other hand, lossless compression maintains a unique pixel location for an element and no averaging of their values occur. The Snow-fall algorithm is a lossless compression algorithm. However, depending upon the memory requirements of a given hardware by convolutional neural network, the size of the pixel frame can be adjusted such that all the elements are represented in the frame, but with lossy compression. Please refer to Supplement File 4 for details about the snow-fall compression algorithm.

### CNN architecture for feature selection and classification

Since class activation maps (CAMs) ^21^ cannot be used for networks that have multiple fully connected layers at the output layer, we used SqueezeNet in this work ^20^. Other nets (e.g. AlexNet, VGG-16, VGG19) can also be used to find CAMs.

The SqueezeNet architecture of DeepInsight-FS has fixed input size (see Figure 3a). It has various hyperparameters such as momentum, L2 regularization, and learning rate. These hyperparameters are tuned on the training set, and the model’s fitness is evaluated on the validation set by employing a Bayesian optimization technique for all the trials. The hyperparameters are selected to minimize the validation error. The test set is never been used in the training or model fitting steps. The chosen CNN hyperparameters produce the optimum performance on the validation set. Description of the parameters is further discussed in Supplement File 1.

**Figure 3:**
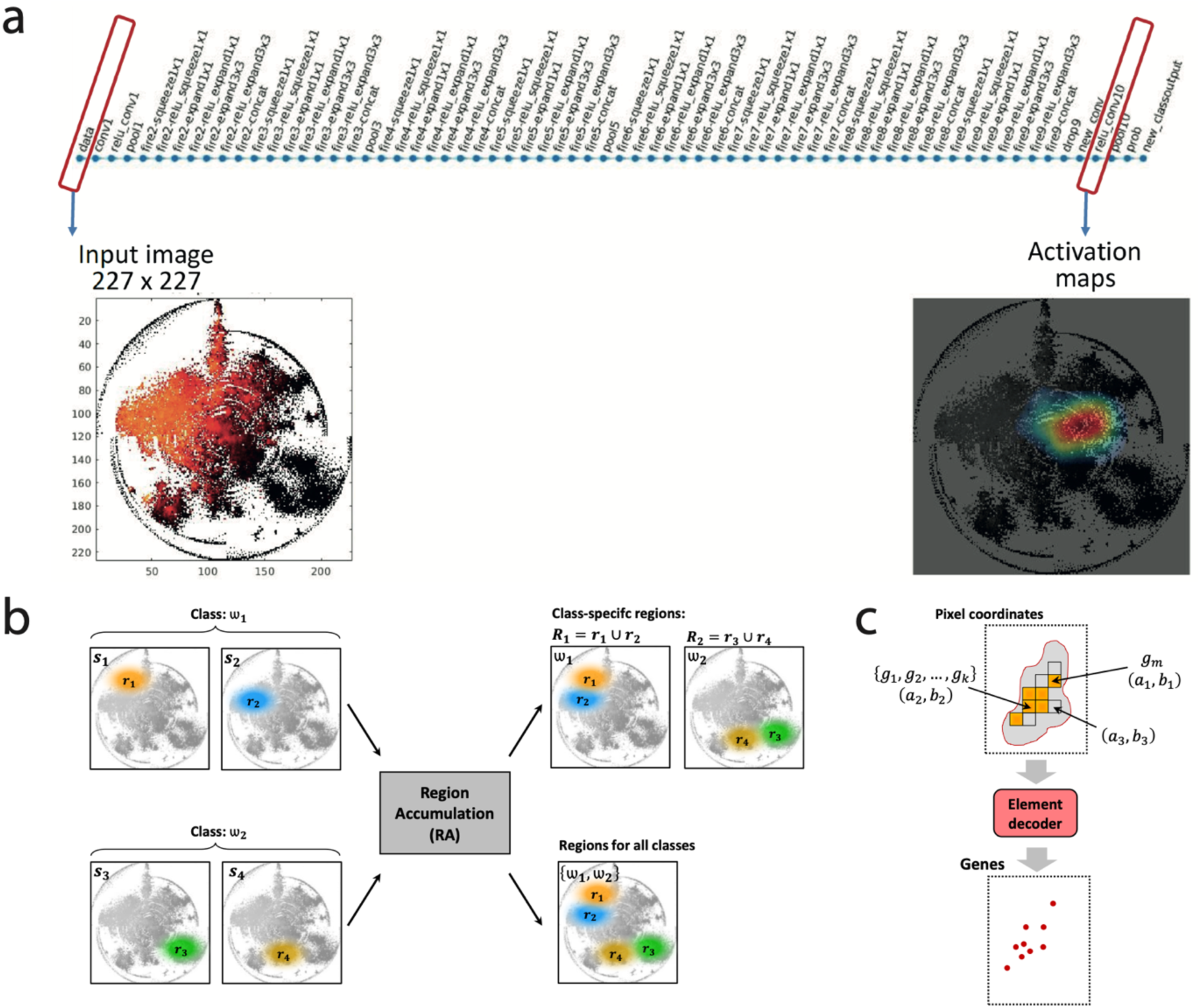
a) SqueezeNet structure with 227 × 227 input image sample from DeepInsight and snow-fall algorithm, and corresponding class activation map at the output ReLu layer. The input image of size *N* × *M* is adjusted to 227 × 227, the image of size of the first convolutional layer of SqueezeNet is *conv1*, and activation maps are retrieved from the last connected ReLu layer (*ReLu_conv10*). After training the CNN model on the optimal hyperparameters, each new sample could be categorized into one of the classes or phenotypes (*new_classoutput*) at the output layer. Here, a sample from RNA-seq data has been first converted to an image by DeepInsight and snow-fall compression algorithm, and then used as an input. At the input, the RNAseq sample can be visualized and a particular region leading to its identification can be analyzed at ReLu layer. The activation map at ReLu layer defines which localities of an image is of interest for decision making process. This activation map has 3 colors in order of preference as red, yellow and blue. The red zone is the most active and blue is the least. b) Region accumulation (RA) step for the DeepInsight-FS method. Two out of four samples belong to a class; i.e., samples {*s*_1_, *s*_2_} are from category ω_1_, and samples {*s*_3_, *s*_4_} are from ω_2_. The active region in each of the samples is depicted as {*r*_1_, *r*_2_, *r*_3_, *r*_4_}. All the four samples are processed via RA step, and it gives 2 outputs. In the first output, integrated regions *R*_*j*_ (where *j* = 1,2), belongs to individual categories are shown; i.e., active regions related to a particular phenotype. In the second output, all the active regions are integrated as *R*, depicting the necessary active zones for classification of two phenotypes. c) Element decoder for the DeepInsight-FS method, the region *R* is supplied to the element decoder. A pixel under *R* could have three possibilities: a pixel representing only one gene *g*_*m*_ at location (*a*_1_, *b*_1_), a pixel representing multiple genes (*g*_1_, *g*_2_, …, *g*_*k*_) at location (*a*_2_, *b*_2_), and a pixel that has no element, e.g. at location (*a*_3_, *b*_3_) (here a pixel value or Base is 1, see Supplement File 1 for further discussion).

### Region accumulation

The region accumulation (RA) step integrates regions of importance. Let a training set with *N* samples be depicted as *S* = {*s*_1_, *s*_2_, …, *s*_*N*_}. Let the activation region corresponding to *N* samples be given as ℋ = {*r*_1_, *r*_2_, …, *r*_*N*_}; i.e., cardinality of ℋ is same as *S*; i.e., |ℋ| = |*S*|. Let *c* be the number of categories (or phenotypes) defined as Ω = {ω_1_, ω_2_, …, ω_*c*_}. Each of the sample will have one of these categories; i.e., *s*_*i*_ *∈* Ω and *r*_*i*_ *∈* Ω, for *i* = 1,2, …, *N*. The overall region of the training data *S* can be evaluated by performing *union operation* of individual regions; i.e. *R* = *r*_1_ ∪ *r*_2_ … ∪ *r*_*N*_ is the integrated region for all samples or for all categories. The region per class is also important to find genes belonging to a particular phenotype. In this case, 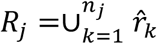, where *n*_*j*_ is the number of samples in the subset represented by ω_*j*_ category, the region 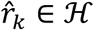 for *k* = 1,2, …, *n*_*j*_ and belongs to a particular class,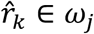. Basically, *R*_*j*_ is the union of all regions depicting a particular class or phenotype. The active region size can vary depending on the threshold value. An illustration is given in Figure 3b.

### Element decoder

The output of RA is processed to the element decoder model. The pixels underneath the selected region are considered for this task. Figure 3c shows the element decoder system. In general, a pixel *p*_*i*_ will have a normalized value 0 ≤ *v*_*i*_, ≤ 1. The decoder will find the argument or index of this pixel *p*_*i*_ located at (*a*_*i*_, *b*_*i*_); i.e., the unique elements or genes, *G*_*i*_, that it contains. If the compression (see the section describing the snow-fall compression algorithm) is lossy then chances are high to get |*G*_*i*_| > 1, and if lossless then |*G*_*i*_| = 1, where | · | is its cardinality. In general, for some pixels |*G*_*j*_| = 1 and for some |*G*_*k*_| > 1 in a given pixel frame (where *j* and *k* are any two pixels).

For a region *R*, a subset of elements or genes will be obtained. It is possible to find a subset of genes for a particular class and also for all the classes. Furthermore, the element decoder can also reveal a subset of genes for a sample. It should be noted that the model can give different subsets of elements for different categories enabling class-dependent findings.

### Reduction of elements in multiple stages

For genomic or transcriptomic data, the number of genes is normally very high and it becomes very difficult to put all genes in a finite image size due to fixed hardware limitations. In this case, it is inevitable to get lossy images; i.e., some image pixels will carry multiple genes in a location. One might wonder, how to perform the selection on those batch genes (where batch gene refers to a set of two or more genes having the same pixel location in the frame). One may also want to reduce the number of selected elements. These issues can be addressed by running DeepInsight-FS in multiple stages. The first stage will find a subset of elements, which can be used as an input for the second stage. Continuing this procedure will reduce the number of elements.

### Experimental setup

We obtained RNA-seq data from the TCGA project. To maintain large enough categories, only 10 types of cancers were considered, namely, TCGA-BRCA, TCGA-COAD, TCGA-HNSC, TCGA-KIRC, TCGA-LGG, TCGA-LUAD, TCGA-LUSC, TCGA-PRAD, TCGA-THCA, and TCGA-UCEC. A total of 6,280 HTSeq-FPKM-UQ expression files were downloaded using the GDC data transfer tool.

From these, 64 files were removed for sharing submitter IDs, resulting in a final total of 6,126 samples. The FPKM-UQ files contain expression for 60,483 genes. In this study, we only used the 19,086 genes classified as protein-coding genes by the HUGO Gene Nomenclature Committee (download date: 2019/11/22).

The dataset was partitioned into 80:10:10 segments of training set, validation set and an independent test set, respectively. The training set is employed to achieve model fitting, whereas the validation set is used to evaluate its fitness. This picks the hyperparameters for which the validation error is small. The independent test set is held aside and applied to provide an unprejudiced evaluation of the final model.

### Performance evaluation

The main aim of this experiment is to show that a subset of imperative elements or genes of non-image data can be selected by CNN with the utilization of DeepInsight-FS method. We have applied enrichment analysis tools like KEGG ^25-27^ and Molecular Signatures Database (MSigDB) ^28,29^ to find out importance of the selected genes to phenotypes by the proposed deep-learning based technique. DeepInsight-FS can find different subset of genes for each of the phenotypes.

We compared our methodology with traditional machine learning techniques. A number of machine learning algorithms exist to find a subset of genes ^30,31^. We applied ANOVA, Lasso ^32^, highly variable genes method ^33^ and logistic regression ^3^ for feature selection, and random forest as a classifier. The feature selection step was applied on the training set. This gives a subset of genes. Since enrichment analysis usually requires a smaller set of genes, the aim is to find roughly ∼1500 genes. Thereafter, hyperparameters of random forest (RF) is tuned using training set on a subset of selected genes, and model fitness is evaluated using the validation set. The classification accuracy (rounded off to its nearest integer) was from the independent test set.

### Evaluation of feature selection capabilities

As outlined above, both DeepInsight-FS and other exemplar algorithms offer some functionality for reducing the number of genes to a more focused subset enriched for the genes used for correctly identifying relevant cancer types. This feature selection is of particular importance in biological data analysis where identification of key genes and underlying mechanisms is usually also part of the overall goal – especially for tasks like identification of clinically-useful sparse diagnostic signatures. To evaluate the utility of DeepInstight-FS from this perspective, we have quantified the enrichment of “gold standard” cancer-specific gene sets and pathways from KEGG and MSigDB (C6 subset). In all instances the enrichment was calculated using Fisher’s exact test and reported p-values were corrected for multiple testing using Benjamini-Hochberg FDR method. Additionally, we report the housekeeping gene counts based on the annotation from The Human Protein Atlas (THPA, http://www.proteinatlas.org) ^34^ and Eisenberg and Levanon 2013 ^35^. Here the assumption is that a more relevant dataset would tend to have fewer housekeeping genes.

### Running the DeepInsight-FS algorithm

Analyzing a large number of elements will cause overlaps in the small pixel-frame and it becomes challenging to perform feature selection. This can cause important elements to be overlooked in the selection process. Therefore, it is useful to perform element reduction to reach a manageable size due to the pixel-frame size and hardware limitations.

The element arrangement step of DeepInsight-FS utilizes t-SNE, which supports various distance measures, *dist*_*j*_. In this study, *dist*_*j*_ are Chebyshev, cosine, correlation, and hamming. A gene set, *G*, processed to DeepInsight-FS with a distance *dist*_*j*_ of t-SNE, gives a gene subset *g*_*j*_. Since four distances are adopted, a union of gene subsets are retrieved, i.e.,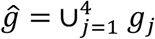. Furthermore, the gene subset ĝ is sent to DeepInsight-FS with hamming distance until a subset of around 1500 genes are obtained (see Figure S1.1 and corresponding discussion in Supplement File 1). Thereafter, gene set and pathway enrichment analyses were performed to evaluate overall relevance of the recovered gene set with respect to current knowledge.

## Supporting information

Video1_SnowFall

## Acknowledgement

The results shown here are in whole or part based upon data generated by the TCGA Research Network: https://www.cancer.gov/tcga. This work was funded by JST CREST Grant Number JPMJCR 1412, Japan; JSPS KAKENHI Grant Numbers JP17H06307, JP17H06299, and JP20H03240; Grant-in-Aid for Scientific Research (JP16H06299) from the Ministry of Education, Culture, Sports, Science and Technology of Japan.

## Author contributions

AS perceived, designed the classification and feature selection models, and wrote the first draft and contributed in the subsequent versions of the manuscript. AL designed the enrichment analysis, improving selection models and contributed in the first draft and thereafter of the manuscript. KAB build further the enrichment analysis, prepared the data, and helped in the manuscript writeup. EV provided art support, build the machine learning tools and contributed in the writeup. TT perceived and contributed in the manuscript writeups. All authors read and approved the manuscript.

## Code Availability

DeepInsight-FS package is available in Matlab to download from link http://emu.src.riken.jp/ or http://www.alok-ai-lab.com/tools.php. Dataset is available from link http://emu.src.riken.jp/DeepInsightFS/dataset1.mat and ReadMe/user manual is available from link https://alok-ai-lab.github.io/deepinsight-fs/

