## Supplement-File_1,2,4 for "DeepInsight-FS: Selecting features for non-image data using convolutional neural network"

### Supplementary File 1
### Feature selection and tuning of DeepInsight-FS parameters

<u>CNN parameter</u>
The SqueezeNet has been used for CNN implementation. The SqueezeNet configuration has a depth of 18. The hyperparameters are obtained by applying the Bayes optimization technique from a range of values. The parameters considered are learning rate, momentum, and l2regularization. Some other parameters are fixed. The maximum objective evaluation and maximum epochs are set to 50 for performing the Bayesian optimization technique. Table S1.1 depicts a summary of the parameters used.

Table S1.1: DeepInsight-FS training parameter options for Bayesian optimization technique

| *Variables* | *Values/range* |
|---|---|
| Net | SqueezeNet |
| Training option | sgdm |
| InitialLearningRate | [1e-5 1e-1] |
| Momentum | [0.8 0.95] |
| L2regularization | [1e-10 1e-2] |
| Max Objectives | 50 |
| MaxTime | 24x60x60 |
| Execution environment | Single-GPU |
| MaxEpochs | 15 |
| Min batch size | 10 |
| Weight learn rate factor | 10 |
| Bias learn rate factor | 10 |
| Image Size | 227 x 227 |

The range of values are applied during the training session, and the best values were selected, which gave the least validation error.

Two types of norms are introduced in DeepInsight [1]. However, in this work, we applied Norm-2.

<u>Dimensionality reduction technique</u>
In many applications the dimensionality of feature space (i.e. the number of features) is very large. It is, therefore, necessary to reduce the dimensionality for better generalization capability, reduced computational complexity, and retrieving information.

The two major streams of dimensionality reduction techniques (DRTs) are feature selection and feature extraction. Feature selection methods retain only a few important features and discards others. On the other hand, feature extraction methods, construct a few features from the large set through their linear (or non-linear) combination. The target or output in this case is usually the addition of weighted inputs (in some cases including biases).

Though, CNN performs feature extraction at its layes, we utilized t-SNE [2] for finding locations of elements or genes. For t-SNE, if the number of elements or data dimensionality is above 5000, Burneshut algorithm is applied for faster processing; otherwise, the exact algorithm is used. To acquire a variety of elements, we utilized various distances of t-SNE such as cosine, correlation, Chebyshev and hamming.

Element versus feature

In this work the term 'element' refers to the raw data, and 'feature' refers to the processed or extracted data. However, the term 'feature' encompasses 'element' since element becomes feature if identify transform is applied, i.e., $feature = identity \times element$.

The term 'element arrangement' in this paper is used because we utilized RNA-seq data. However, this term can be considered identical to 'feature arrangement'.

Feature mapping

After determining the element locations using the training set, the mapping of elements is carried on to these locations. In a situation where more than one elements occupy the same spot, then their averaged values are utilized; for instance, if locations of $g_i, g_j$ and $g_k$ are the same $(r, c)$ then $(g_i + g_j + g_k)/3$ will be mapped on this location. In this case, lossy compression will occur. Only the training set is used to find element locations. The validation and test sets used the same locations obtained from the training set. In regards to the empty pixels; i.e., the pixel locations that do not have any elements are referred to as Base, and its value is fixed as 1.

Feature selection using DeepInsight-FS

The RNA-seq data was pre-filtered to reduce to 19086 protein-coding genes. In the training set, several genes have zero expression values, and such 73 genes have been removed. This gives 19013 genes. After that, DeepInsight-FS was used to produce around 1500 genes for enrichment analysis. As discussed under section 'Running the DeepInsight-FS algorithm' of the manuscript, DeepInsight-FS run in multiple stages to acquire the desired number of elements. Figure S1.1 shows an overall framework of DeepInsight-FS with various distances used.

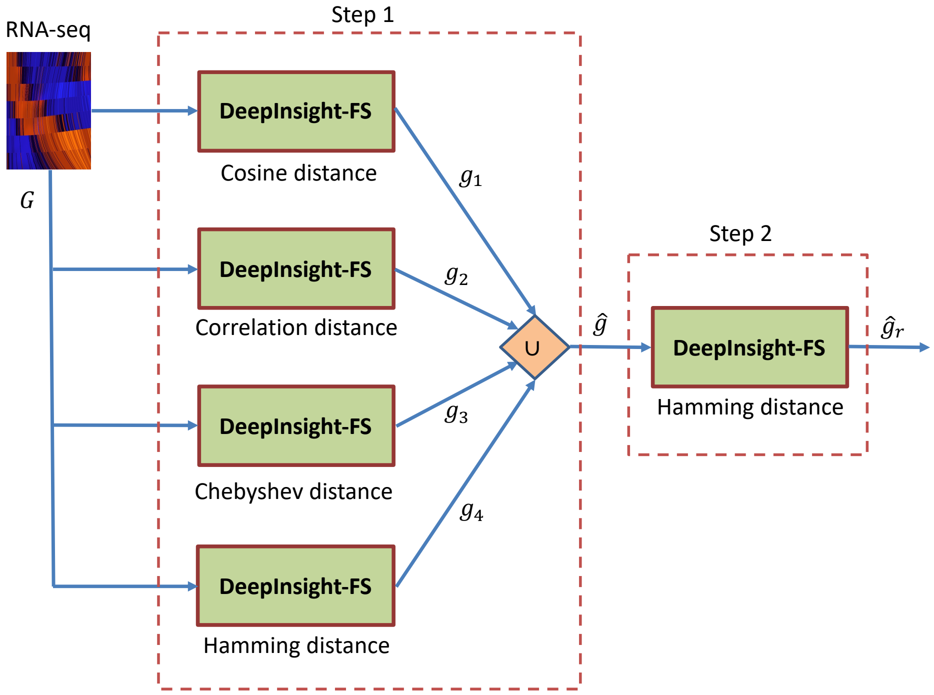


Figure S1.1: An overall framework of DeepInsight-FS execution scheme.

The RNA-seq data, with 19,013 genes, is processed to DeepInsight-FS using different distances at *step 1*. As a result, gene subsets $g_1$, $g_2$, $g_3$ and $g_4$ are found. The union of these four subsets is $\hat{g}$, which is then further processed to DeepInsight-FS with hamming distance at *step 2*, and obtain a final subset $\hat{g}_r$. The details of hyperparameters selected for each run and the number of genes at *step 1* are listed in Table S1.2, and for *step 2* are listed in Table S1.3. The union of genes, $\hat{g}_r$, retrieved from *Step 1* is 5228. Since the gene size is much larger than the desired genes (~1500), *step 2* is conducted to reduce the size further. For *step 2*, the input gene size is 5228 and the output is 1806. The classification accuracy on the independent test set with the selected gene subset is 98%.

Table S1.2: Selected hyperparameters and number of genes selected during each stage of DeepInsight-FS in *Step 1*.

1. *DeepInsight-FS with cosine distance*

| | Stage1 | Stage2 | Stage3 |
|---|---|---|---|
| **#Genes** | 7635 | 4851 | 2381 |
| **Val error** | 0.017889 | 0.0143 | 0.017889 |
| **InitialLearningRate** | 0.0001416 | 1.0993E-04 | 3.85E-04 |
| **Momentum** | 0.87056 | 0.9316 | 0.8021 |
| **L2Regularization** | 0.0067864 | 2.2359E-05 | 2.81E-09 |
| **CAM Threshold** | 0.6 | 0.6 | 0.6 |
| **ImageSize** | 227 x 227 | 227 x 277 | 227 x 227 |

2. *DeepInsight-FS with correlation distance*

| | Stage1 | Stage2 |
|---|---|---|
| **#Genes** | 8361 | 1603 |
| **Val error** | 0.0143 | 0.0179 |
| **InitialLearningRate** | 2.51E-05 | 2.5033E-04 |
| **Momentum** | 0.943679 | 0.931836 |
| **L2Regularization** | 4.99E-09 | 3.0133E-10 |
| **CAM Threshold** | 0.6 | 0.6 |
| **ImageSize** | 227 x 227 | 227 x 277 |

3. *DeepInsight-FS with Chebyshev distance*

| | Stage1 | Stage2 | Stage3 | Stage4 |
|---|---|---|---|---|
| **#Genes** | 5952 | 3415 | 2210 | 1460 |
| **Val error** | 0.0125 | 0.0161 | 0.0161 | 0.0215 |
| **InitialLearningRate** | 6.02E-05 | 3.3292E-05 | 2.43E-05 | 0.00021184 |
| **Momentum** | 0.918806 | 0.895583 | 0.942427 | 0.867717 |
| **L2Regularization** | 7.93E-03 | 1.1713E-10 | 2.10E-04 | 9.32E-07 |
| **CAM Threshold** | 0.6 | 0.6 | 0.6 | 0.6 |
| **ImageSize** | 227 x 227 | 227 x 277 | 161 x 158 | 149 x 133 |

4. DeepInsight-FS with hamming distance

| | Stage1 | Stage2 |
|---|---|---|
| **#Genes** | 5188 | 2469 |
| **Val error** | 0.0161 | 0.0143 |
| **InitialLearningRate** | 2.93E-04 | 9.2110E-07 |
| **Momentum** | 0.805893 | 0.934098 |
| **L2Regularization** | 2.64E-03 | 9.5067E-03 |
| **CAM Threshold** | 0.6 | 0.6 |
| **ImageSize** | 227 x 227 | 227 x 227 |

Table S1.3: Selected hyperparameters and number of genes selected in *step 2*.

| | Stage1 |
|---|---|
| **#Genes** | 1806 |
| **Val error** | 0.0125 |
| **InitialLearningRate** | 4.07E-04 |
| **Momentum** | 0.923147 |
| **L2Regularization** | 1.97E-09 |
| **CAM Threshold** | 0.45 |
| **ImageSize** | 227 x 227 |

Figure S1.2 shows an input RNA-seq sample having 5228 genes (left) and its activations at the output (right). The preferential order of activation is from red to yellow to blue; i.e., the genes under the red activation zone is the most desirable. We have only considered genes corresponding to the red activation and discarded the other regions such as yellow, blue and without activation. After processing all the training set, we get 1806 genes in total for all the 10 cancer types.

DeepInsight-FS gives different gene subsets for different phenotypes. This is depicted in Figure S1.3. In this figure, class 1 to class 10 refer to cancer types TCGA-BRCA, TCGA-COAD, TCGA-HNSC, TCGA-KIRC, TCGA-LGG, TCGA-LUAD, TCGA-LUSC, TCGA-PRAD, TCGA-THCA, and TCGA-UCEC, respectively. For these cancer types, the DeepInsight-FS method selects the following number of genes (listed in the same order of cancer types as mentioned above): 1444, 1147, 1277, 1341, 936, 1135, 1167, 1210, 1036 and 1294, respectively. For many cancer types, the same or common genes appeared, and the union of genes belonging to 10 cancer types is 1806.

It can be observed from Figure S1.3 that most of the genes to represent cancers are same. However, some genes are different, which helps in identifying different cancers. Therefore, the DeepInsight-FS method finds genes which are representative and at the same time different for each cancer type.

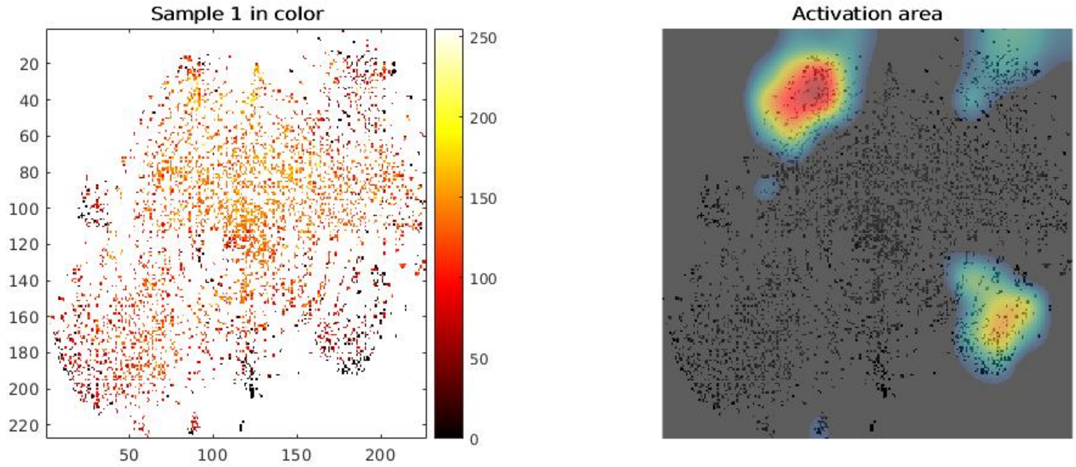


Figure S1.2: DeepInsight-FS: a converted image sample having 5228 genes (left) and its corresponding activations (right).

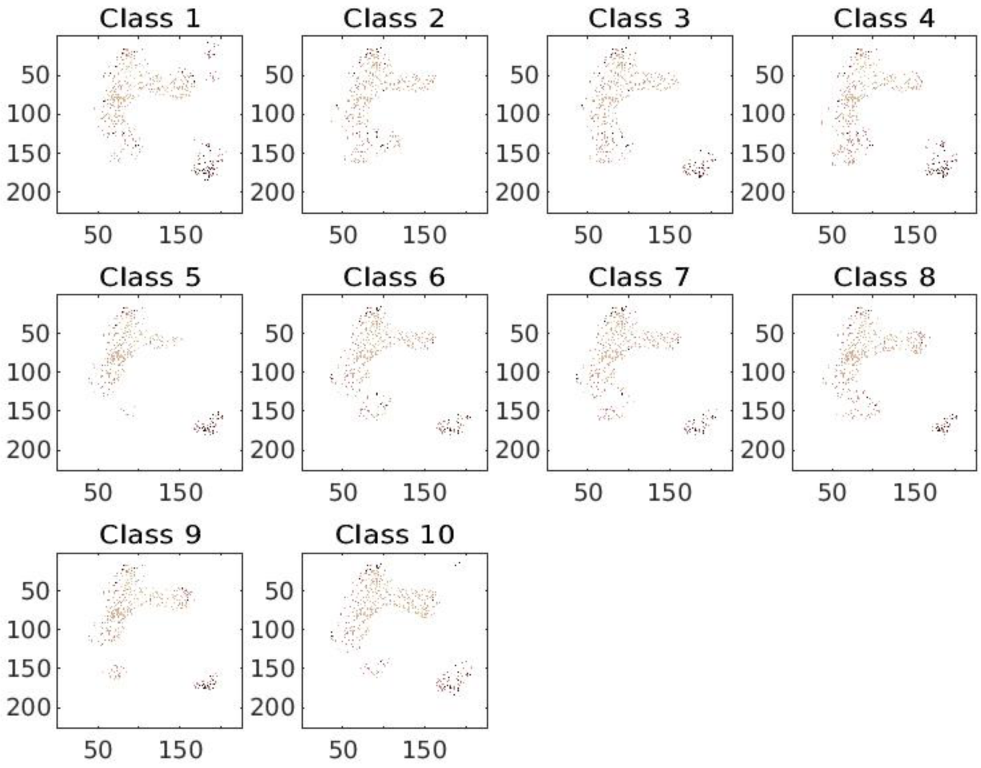


Figure S1.3: Selected gene subsets of each 10 cancer types.

### Supplementary File 2
### Enrichment analysis

Figure S2.1 illustrations gene set overlap by DeepInsight-FS and logistic regression techniques. The gene overlap is highest between the same class. Considerable overlap between adjacent columns of DeepInsight-FS can be seen for DeepInsight-FS, depicting similar genes are used to identify different cancer types.

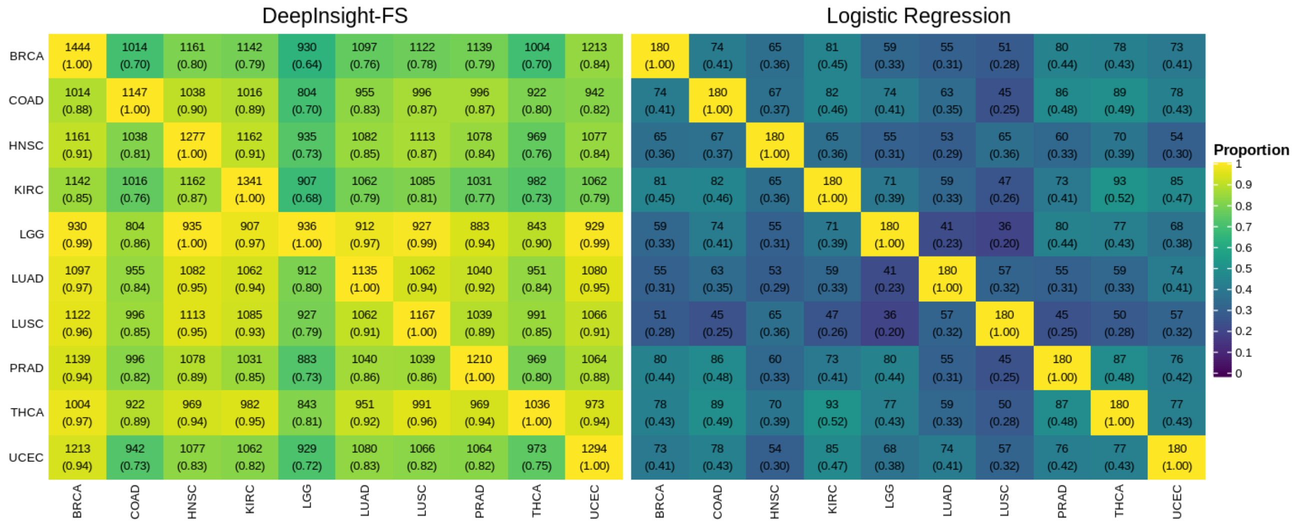


Figure S2.1: Gene set overlap by DeepInsight-FS and logistic regression

Figure S2.2 illustrates the performance evaluation of all the methods studied in this work in terms of the number of significant enrichment pathways. We performed KEGG and MSigDB analyses on the gene sets obtained from different algorithms. It can be seen that ANOVA, Lasso and high-variance genes (HVG) methods did not obtain unique gene sets for individual cancer types. Nonetheless, logistic regression and DeepInsight-FS perform better than the other 3 methods.

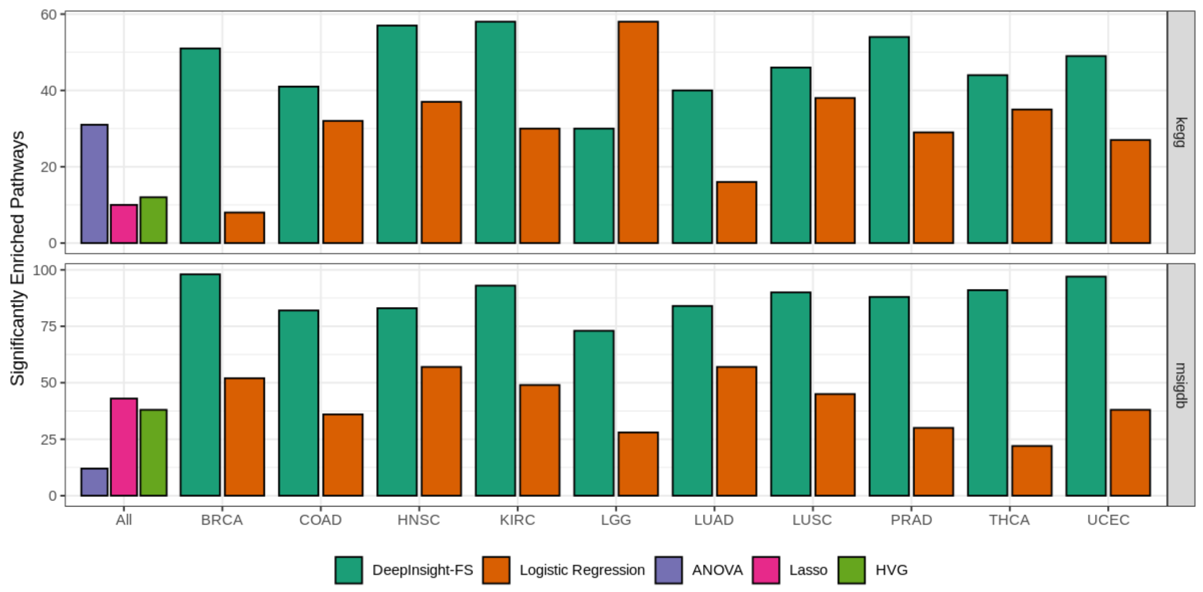


Figure S2.2: KEGG and MSigDB analyses on ANOVA, Lasso, high-variance genes, logistic regression and DeepInsight-FS.

Figure S2.3a shows overlap of gene subsets between several methods studied in this paper. For an instance, ANOVA and DeepInsight-FS (DI-FS) has 143 genes in common, and there are no common genes between ANOVA and HVG. Lasso and DI-FS have 205 common genes which is the highest among any two methods. Moreover, Figure S2.3b depicts gene annotation of all the methods. Yellow region shows the selected genes for a particular method (in column), and purple region shows genes non-selected. It also shows HouseKeeping genes. For example, HVG has the least number of HouseKeeping genes compared to other methods tested.

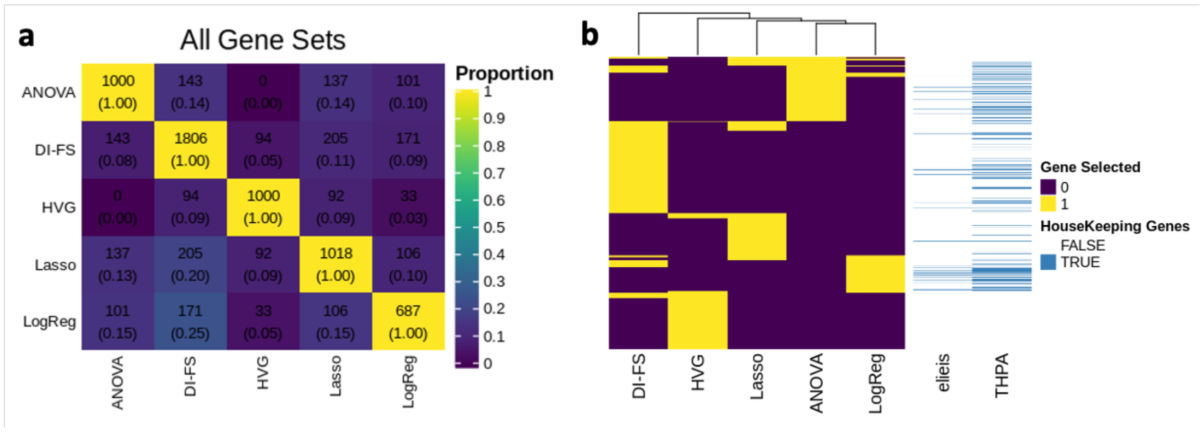


Figure S2.3: a) Overlap of gene subsets between all the methods; and b) gene annotation for all the methods (where DI-FS is DeepInsight-FS).

### Supplementary File 4
### Snow-fall compression algorithm

For an illustration, consider Figure S4.1a, where a $6 \times 6$ pixel frame represents 5 elements or $x \in \mathbb{R}^5$. The pixel frame has been divided into 4 quadrants. If the distance of an element $i$ from the center C is depicted as $d_i$ then according to Figure S4.1, $x_3$, is the closest from the center C (shaded as grey), and $x_1$ is the farthest. Since the distance $d_3$ is lowest, element $x_3$ will first move or fall towards the center, followed by $x_2$, $x_5$, $x_4$, and $x_1$. The falling towards the center $C$ will continue until the element reaches $C$ or hit by some other elements on its path. This will free up space and as a result, it is possible to represent more elements in a given pixel frame.

After moving all the elements towards the center, the frame will appear as shown in Figure S4.1b (as blue dots). It can be seen that $x_3$ captured the center position, and other elements made free fall to squeeze towards the center of the frame. Now, instead of having a frame size of $6 \times 6$, we can manage with the frame size of $3 \times 4$. This frees up more space and new elements can be added (as shown in Figure S4.1b as red dots) or in other words higher dimensional data can be considered for the same pixel frame. More elements e.g. $x_6, x_7$ and $x_8$, are added to the same frame in Figure S4.1b after falling of elements towards $C$.

In this algorithm, the elements fall towards the center of the frame, and hence we call it a snow-fall compression algorithm. In the next section, we discuss the mathematics of the snow-fall algorithm.

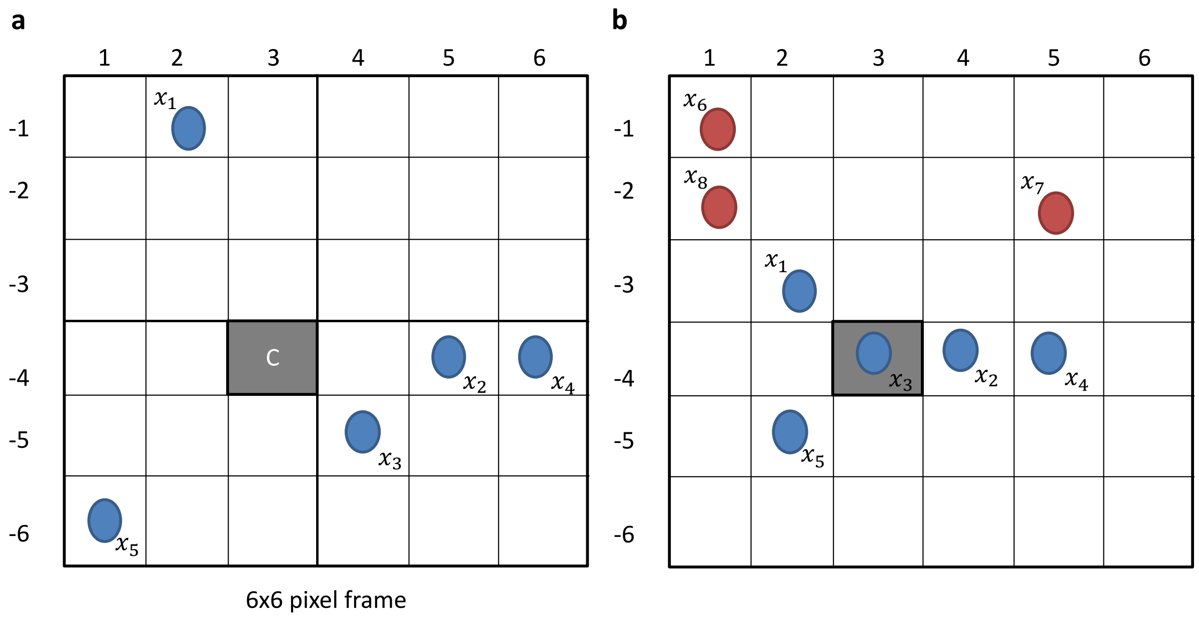


**Figure S4.1**: a) An illustration of 5 elements $x_i$ (for $i = 1,2,\ldots,5$) in a 6x6 pixel frame (note rows are depicted in negatives so that cartesian coordinates and pixel coordinates can be represented simultaneously). b) Squeezing all the elements towards the center, and adding new elements or features after compression (red dots).

***Basis of Snow-Fall Algorithm***

Let $U = \{u_1, u_2, \ldots, u_d\}$ be the vectors in 2D pixel frame. The first step finds a quadrant where a vector or point $u = (x_u, y_u) \in U$. In Figure S4.2a, the four quadrants are shown and conditions are given for when a point $u$ will come under one of these 4 quadrants. The center point is depicted as $u_c = (x_c, y_c)$ and the edge point is $u_p = (x_p, y_p)$.

Given points $u$, $u_c$ and $u_p$, the quadrant conditions are specified as

1) Q1 if $x_p \le x_c$ and $y_c \le y_p$, find point $u$ where

$x_p \leq x_u \leq x_c$, and
$y_c \leq y_u \leq y_p$

2) Q2 if $x_p \leq x_c$ and $y_p < y_c$, find point $u$ where
$x_p \leq x_u \leq x_c$, and
$y_p \leq y_u < y_c$
3) Q3 if $x_p > x_c$ and $y_c \geq y_p$, find point $u$ where
$x_p \geq x_u > x_c$, and
$y_c \geq y_u > y_p$
4) Q4 if $x_p > x_c$ and $y_p > y_c$, find point $u$ where
$x_p \geq x_u > x_c$, and
$y_p \geq y_u > y_c$

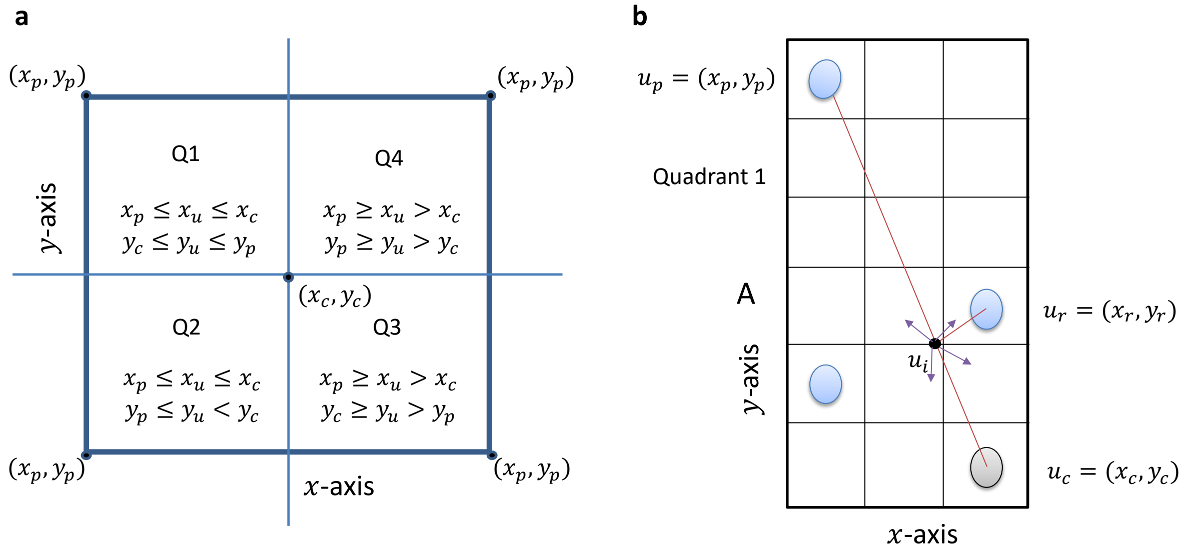


**Figure S4.2**: a) Quadrant analysis to locate a point $(x_u, y_u)$ (note the quadrant numbers are selected as per convenience and it may not be identical to angular quadrants). The frame is adjusted such that no point has negative values. The edge point is depicted by $(x_p, y_p)$ and center as $(x_c, y_c)$ b) An illustration of points under subarea A, where $u_r = (x_r, y_r)$ is the closest point to $u_p = (x_p, y_p)$, and $u_i$ is the intersection point between the two perpendicular lines.

In order to develop the model, let us consider quadrant 1 (Q1) as depicted in Figure S4.2b. Let $A$ be the subarea in Q1, $u_c$ be the center point, $u_p$ be the edge point of $A$, and $u_r$ be the closest point to $u_p$ given subarea $A$.

In $A$, equation of a line emanating from $u_p$ and passing at the center can be depicted as

$$y = mx + (y_c - mx_c) \quad (1)$$

Assuming no points in $A$ exit other than $u_p$, then

Case1) No point at $u_c = (x_c, y_c)$ exists and therefore $u_p = (x_p, y_p)$ will fall at location of $u_c$, i.e.,
$u_c \leftarrow u_p$.

Case 2) If a point $u_c$ exist at the center, then
If $x_p < x_c$, then
$x'_p = x_c - 0.5$, where $x'_p$ is a new location of $x_p$, and from Eq (1)
$y'_p = [mx'_p + (y_c - mx_c)]$, where $[\cdot]$ is either *floor* or *ceil* of a numerical value.

If $x_p > x_c$, then

$x'_p = x_c + 0.5$, and from Eq (1)

$y'_p = [mx'_p + (y_c - mx_c)]$

If other points also exit as depicted in Figure S4.2b, then the model will search closest distance from $u_p$ to all $u \in A$, i.e.,

$$r = \arg\min_{u_j \in A}(u_p, u_j) \quad (2)$$

This will return point $u_r$, a point closest to $u_p$.

Now a perpendicular line to Eq (1) and passing $u_r$ can be given as

$$y = -\frac{1}{m}x + (y_r + \frac{1}{m}x_r) \quad (3)$$

The intersection point $(x_i, y_i)$ of Eq (1) and Eq (3) can be found by substituting Eq (3) in Eq (1) (see intersecting point as a black dot in **Figure S4.**Figure S4.2b), we get

$$x_i = \frac{\left(y_r + \frac{1}{m}x_r\right) - (y_c - mx_c)}{m + \frac{1}{m}} \quad (4)$$

Substituting $x_i$ in Eq (1), we get

$$y_i = \frac{m\left(\left(y_r + \frac{1}{m}x_r\right) - (y_c - mx_c)\right)}{m + \frac{1}{m}} \quad (5)$$

Therefore, in this case, the new location for $u_p = (x_p, y_p)$ would be $u'_p = u_i = (x_i, y_i)$.

Note that these $x_i$ and $y_i$ are in Cartesian coordinates, and it can move to at most 4 locations in pixel coordinates (shown by arrow in Figure S4.2b).

One way of finding the location for intersection point in the pixel coordinate is to compute distance $d_{pi}$ between $u_p$ and $u_i$ as

$$d_{pi} = \sqrt{\left(x_p - [x_i]\right)^2 + \left(y_p - [y_i]\right)^2} \leq d_{pr} \quad (6)$$

Where [·] is either *floor* or *ceil*, and $d_{pr}$ is the distance between $u_p$ and $u_r$; i.e.,

| | | | | | |
|---|---|---|---|---|---|
| $[x_i]$ | = | $floor(x_i)$ | $floor(x_i)$ | $ceil(x_i)$ | $ceil(x_i)$ |
| $[y_i]$ | = | $floor(y_i)$ | $ceil(y_i)$ | $floor(y_i)$ | $ceil(y_i)$ (7) |
| (distance) | | $d_{ff}$ | $d_{fc}$ | $d_{cf}$ | $d_{cc}$ |

This gives 4 conditions, and the best condition would be $d_{pi} < d_{pr}$. Suppose $d_{ff} < d_{fc} < d_{cf} < d_{pr} < d_{cc}$, then select condition $cf$ and accordingly select the new location of $u_p$; i.e., $u'_p = (ceil(x_i), floor(y_i))$.

Another way of finding the location can be considered by using center as a reference point. Let $d_{ci}$ be the distance between the center and the intersection point, we have

$$d_{ci} = \sqrt{(x_c - [x_i])^2 + (y_c - [y_i]^2)} \quad (8)$$

In similar way as Eq (7), [·] can be either *floor* or *ceil*, giving 4 conditions. The nearest point towards the center would be

$$(c', i') = \arg \min_{ff,fc,cf,cc} d_{ci} \tag{9}$$

These steps have to be taken for other points, making this algorithm an iterative one until all the points fall.

***Intersecting points when $m = 0$ and $m = \infty$***

Next, we see the intersecting point (Eq (4) and Eq (5)) in extreme condition; i.e., when $m = \infty$ and $m = 0$.

if $m = \infty$ then from Eq (4)

$$x_i = \frac{\left(y_r + \frac{1}{m}x_r\right) - (y_c - mx_c)}{m + \frac{1}{m}} = \frac{\infty}{\infty} \text{ (intermediate terms)}$$

$$x_i = \lim_{m \to \infty} \frac{y_r}{m + \frac{1}{m}} + \frac{x_r}{m^2 + 1} - \frac{y_c}{m + \frac{1}{m}} + \frac{x_c}{1 + \frac{1}{m^2}}$$

$$x_i = x_c$$

Similarly, from Eq (5)

$$y_i = \frac{m\left(\left(y_r + \frac{1}{m}x_r\right) - (y_c - mx_c)\right)}{m + \frac{1}{m}} + (y_c - mx_c) = \frac{\infty}{\infty}$$

$$y_i = \lim_{m \to \infty} \frac{y_r}{1 + \frac{1}{m^2}} + \frac{x_r}{m + \frac{1}{m}} + \frac{y_c}{m^2 + 1} - \frac{mx_c}{m^2 + 1}$$

$$y_i = y_r + 0 + 0 - \lim_{m \to \infty} \frac{x_c}{2m} \quad \textit{(from L'Hopital's Rule)}$$

$$y_i = y_r$$

If $m = 0$ then $x_i = x_r$ and $y_i = y_c$.

***Lossless and Lossy Compression***

The snow-fall algorithm gives an optimum pixel frame of size $p \times q$. The compression would be lossless if the desired frame size $m \times n$ is greater than $p \times q$ ($p \leq m$ and $q \leq n$). The compression can be lossy if $p > m$ and $q > n$. The lossy compression can be attained by adjusting the horizontal and vertical sizes of the pixel frame. In this case, it is possible that more than 2 elements can attain one location giving the average value of elements at this particular location. For an instance, if $k$ elements $\{g_1, g_2, \dots, g_k\}$ having a common location then average value $\frac{1}{k}\sum_{i=1}^{k} g_i$ will be mapped at this location.

An illustration of a pixel-frame of size $100 \times 100$ extracted by DeepInsight is shown in Figure S4.3a. Its lossless form by snow-fall algorithm of size $15 \times 14$ is depicted in Figure S4.3b, and its lossy compression form of size $8 \times 7$ is given in Figure S4.3c. A video illustration (
Video 1) can be accessed as *SnowFall_Video.avi* (note the speed in the video has been reduced for illustration purpose). In the case of lossy compression, multiple elements or genes occupy a common location.

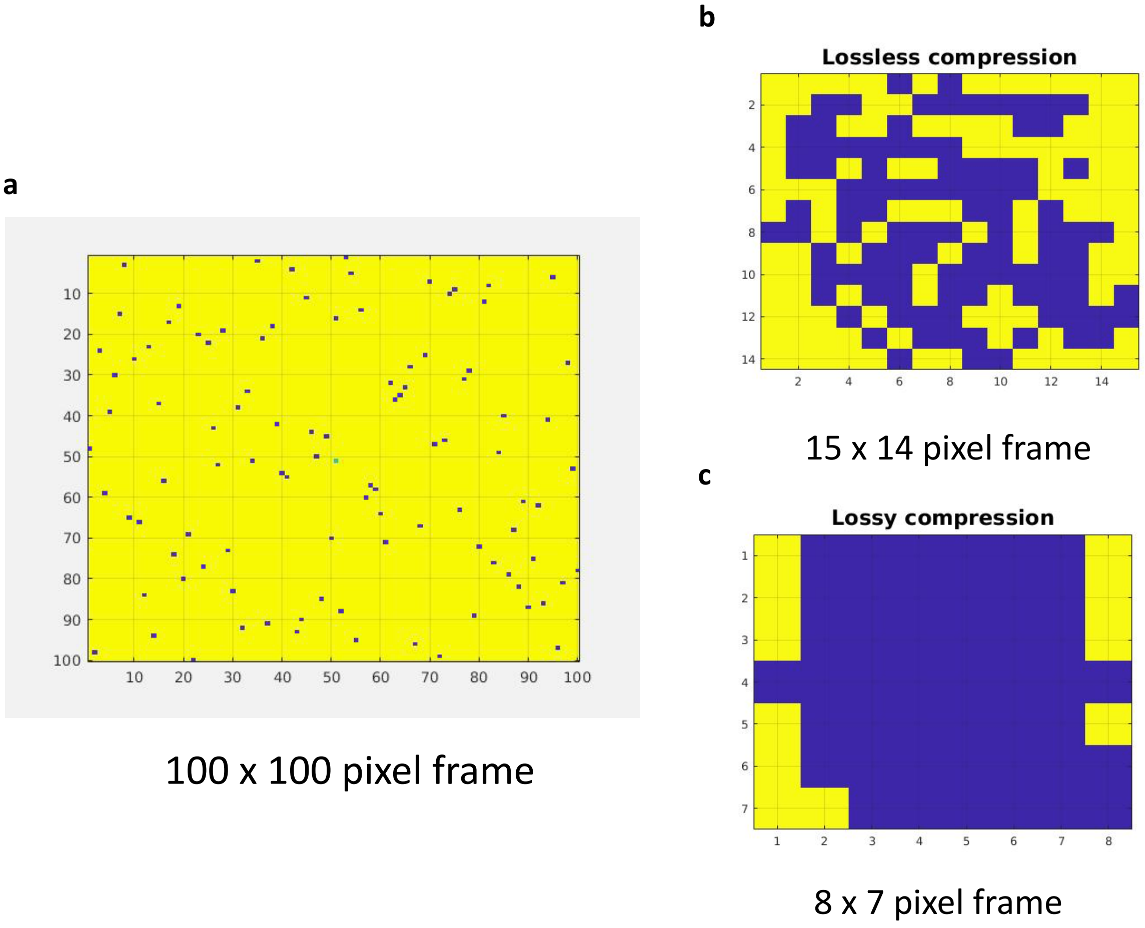


**Figure S4.3**: An illustration of the snow-fall algorithm a) 100 × 100 pixel frame, b) lossless snow-fall compression, and c) lossy snow-fall compression.

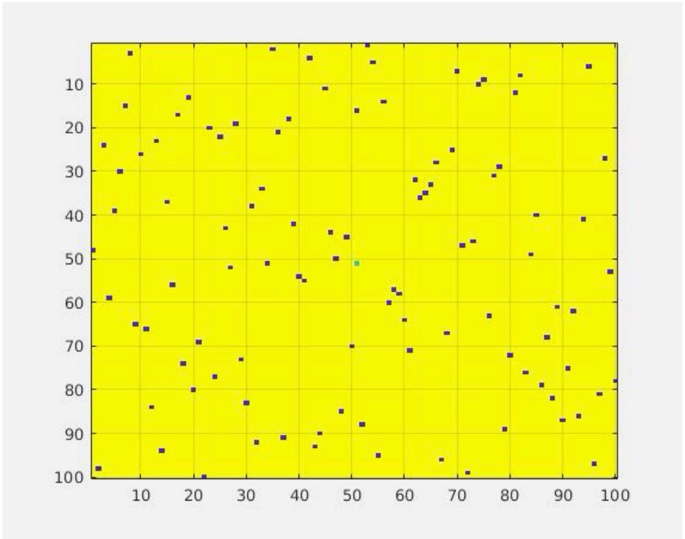

*Video 1: A run of snow-fall compression algorithm.*
